# Sustained hippocampal theta-oscillations reflect experience-dependent learning in backward temporal order memory retrieval

**DOI:** 10.1101/2023.07.10.548388

**Authors:** Hongjie Jiang, Jing Cai, Diogo Santos-Pata, Lei Shi, Xuanlong Zhu, Jiaona Tong, Yudian Cai, Chenyang Li, Rui Wang, Jia Yin, Shaomin Zhang, Sze Chai Kwok

**Affiliations:** The Second Affiliated Hospital of Zhejiang University, School of Medicine, Department of Neurosurgery, Hangzhou, China; Phylo-Cognition Laboratory, Division of Natural and Applied Sciences, Data Science Research Center, Duke Kunshan University, Duke Institute for Brain Sciences, Kunshan, Jiangsu, China; Department of Neurosurgery, China Medical University, The First People’s Hospital of Kunshan, Suzhou, Jiangsu, China; Key Laboratory of Biomedical Engineering of Ministry of Education, Qiushi Academy for Advanced Studies, Zhejiang Provincial Key Laboratory of Cardio-Cerebral Vascular Detection Technology and Medicinal Effectiveness Appraisal, Zhejiang University, Hangzhou, China; Shanghai Key Laboratory of Brain Functional Genomics, Key Laboratory of Brain Functional Genomics (Ministry of Education), Affiliated Mental Health Center (ECNU), School of Psychology and Cognitive Science, East China Normal University, Shanghai, China; Department of Neurosurgery, Shanghai Tenth People’s Hospital, Tongji University, No. 301 Yanchang Road, Shanghai, 200072, China; Shanghai Key Laboratory of Magnetic Resonance, East China Normal University, Shanghai, China; Shanghai Changning Mental Health Center, Shanghai, China; Clinical Research Center for Neurological Diseases of Zhejiang Province, Hangzhou, China

**Keywords:** hippocampus, theta oscillations, episodic memory, intracranial EEG, learning, p-episodes, temporal order judgment

## Abstract

Navigating within our neighborhood, learning a set of concepts, or memorizing a story, requires remembering the relationship between individual items that are presented sequentially. Theta activity in the mammalian hippocampus has been related to the encoding and recall of relational structures embedding episodic memories. However, how theta oscillations are involved in retrieving temporal order information in opposing directionality (forward vs backward) has not been characterized. Here, using intracranial recordings from 10 human epileptic patients of both genders with hippocampal electrodes, we tested the patients with a temporal order memory task in which they learned the spatial relationship among individual items arranged along a circular track and were tested on both forward-cued and backward-cued retrieval conditions. We found that sustained high-power oscillatory events in the hippocampal theta (2-8 Hz) band, as quantified by P_episode_ rate, were higher for the backward conditions during the later stage but not in the earlier stage. The theta P_episode_ results are consistent with the behavioral memory performance. In contrast, we observed a stronger effect of forward than backward retrieval for the gamma (30-70 Hz) P_episode_ rate irrespective of stages. Our results revealed differential roles of theta vs. gamma oscillations in the retrieval of temporal order and how theta oscillations are specifically implicated in the learning process for efficient retrieval of temporal order memories under opposing directionality.

**Significance statement:** While the hippocampus is critical to link events into unitary episodes, the effect of repeated experiences, or learning, on these processes is not entirely clear. We discovered that hippocampal theta oscillation in humans is modulated by repeated experiences, which in turn increases the efficacy of backward-cued memory retrieval of temporal order. This study revealed an important physiological signature characterizing the role of experiences and learning in bidirectional temporal memory retrieval.

**Journal section:** Behavioral/Cognitive

## Introduction

The hippocampus is involved in binding items with their spatiotemporal contexts and encoding sequential order information (Squire, 1992; Eichenbaum, 2000). Following the discovery of place cells, Mehta et al. (1997) found that the receptive field of a place cell acquires a negatively skewed shape through experience, such that the firing rate is initially low as the animal enters the field but will increase as it exits the arena. Such experience dependent effects are consistent with place field changes observed during the phase precession effect, where the animal traverses a cell’s place field the phase position in which the place cell fires within a theta cycle advances (O’Keefe & Recce, 1993; Skaggs et al, 1996; Tsodyks et al, 1996). In a later study, Mehta et al. (2000) proposed that the skewness acquired through experience may reflect the directional synaptic strengthening caused by time lag between the pre- and postsynaptic spikes from CA3 to CA1.

In temporal order memory tasks, the encoding of a stimulus could be linked with its “temporal context” in a directional manner, such that this stimulus is associated with those stimuli immediately preceding it more strongly than those following it. By the Temporal Context Model (Howard & Kahana, 2002), retrieved context is an inherently asymmetric retrieval cue and would lead to a widespread advantage for forward recalls compared to backward recalls (Michelmann et al, 2019; Yang et al, 2013; Kahana & Caplan, 2002). Ample behavioral evidence also confirmed that forward and backward recalls implicate different retrieval processes (Li & Lewandowsky, 1995; Chan, Ross, Earle *et al*, 2009; Liu, Chan & Caplan, 2014; Liu & Caplan, 2022). However, how retrieval directionality in memory tests might be modulated by the effect of learning, or familiarization of one’s environment, remains unknown.

Putting the aforementioned experience-dependent phenomena to memory retrieval of temporal order, it is possible that the experience-dependent skewness property applies on a temporal dimension. Through experience, hippocampal oscillations facilitate temporal sequence learning by compressing and replaying the temporal order of events by transforming rate coding to temporal coding (Mehta, Lee & Wilson, 2002). In the human literature, theta oscillations in the hippocampal formation are analogously involved in spatial navigation (Ekstorm et al, 2003, 2005; Liu et al, 2023) and memories (Solomon et al, 2019; Lega, Jacobs & Kahana, 2012), akin to the function of rodent place cells. Indeed, human MTL neurons also exhibit a similar form of phase precession during visual learning (Reddy et al, 2021) and item-context binding (Zheng et al, 2022). These suggest that the theta oscillations in the human hippocampus provide a proxy for us to examine the effects of experience-dependent property on memory retrieval. However, it remains to be elucidated how such experience-dependence could affect the efficacy of retrieval directionality (forward vs backward) during memory judgments of relative temporal order.

In the current study, we recorded hippocampal local field potential activity via intracranial EEG in epileptic patients while they performed a temporal order memory test. Participants viewed through a circular track in virtual reality and then made a relative order memory judgment between two items with respect to a sample. We manipulated the extent of how the participants experienced the virtual reality environment by dividing the experiment into three stages and compared cued judgments of temporal order for forward versus backward sequences. Neurally, we used sustained oscillations in the hippocampus - the P_episodes_ (Caplan et al, 2001, 2003; Whitten et al., 2011) - to verify our hypotheses. As implied by experience-dependent, negatively skewed receptive fields (Mehta et al, 1997, 2000), theta oscillations in the hippocampus should disproportionally reflect higher activation for backward-cued retrieval as the experiment progressed. These effects will be accompanied with a change in memory performance that backward-cued retrieval progressively becomes more accurate. We confirmed the relationship between experiences and directionality during memory test is theta-specific by including an analysis on P_episode_ rate in the gamma range as a control comparison.

## Materials and Methods

### Participant details

Ten medically refractory epilepsy patients were recruited while they were hospitalized for pre-surgical diagnosis at the Department of Neurosurgery, the Second Affiliated Hospital of Zhejiang University. The patients had been surgically implanted with depth electrodes for diagnostic screening, preceding the surgical treatment (Table 1). All patients underwent neuropsychological evaluation and had a normal or corrected-to-normal vision. They were informed about the procedure and provided written consent before participating in the experiment. The study was approved by the Human Research Ethics Committee of the Second Affiliated Hospital of Zhejiang University School of Medicine (IR202001335).

**Table 1.** Patients’ demographic profile and clinical conditions.

| Sub ID | Age | Gender | Implantation sites | Seizure onset | Resection | Psychological deficits | Pathological conditions |
| --- | --- | --- | --- | --- | --- | --- | --- |
| 01_YG | 23 | M | bilateral temporal lobe, right frontal lobe & insula | bilateral temporal lobe & right precentral gyrus | none | mild hypomnesia | none |
| 02_JY | 25 | M | bilateral frontal lobe, right temporal lobe & insula | right temporal lobe | right Anterior Temporal Lobectomy | moderate hypomnesia, anxiety, phobia | FCD Type IIa (Focal Cortical Dysplasia) |
| 03_LYY | 7 | F | bilateral occipital lobe, left temporal lobe | left occipital lobe | left occipital epileptogenic focus | visual illusions during seizure period | FCD Type I (Focal Cortical Dysplasia) |
| 04_YY | 18 | M | right frontal, temporal, parietal lobe & insula | right temporal & inferior parietal lobe, cingulate gyrus, insula | none | mild hypomnesia | none |
| 05_CLC | 20 | M | left frontal, temporal & parietal lobe | left frontal lobe | left frontal epileptogenic focus | motor dysfunction in right upper limb during seizure period | Gliocyte proliferation |
| 06_SCM | 35 | F | bilateral temporal lobe, left frontal lobe & insula | left temporal lobe | left Anterior Temporal Lobectomy | moderate hypomnesia, anxiety, phobia | Gliocyte proliferation |
| 07_LZ | 24 | M | left temporal lobe, cingulate gyrus, insula, parietal lobe & occipital lobe | left anterior cingulate gyrus & anterior insula | none | mild hypomnesia | none |
| 08_ZLX | 14 | M | bilateral temporal lobe, left frontal lobe, insula & parietal lobe | left temporal lobe | left Anterior Temporal Lobectomy | moderate hypomnesia | Gliocyte proliferation |
| 09_WCF | 34 | F | right frontal lobe, temporal lobe, cingulate gyrus & insula | right frontal lobe & cingulate gyrus | none | mild phobia | none |
| 10_LGD | 29 | M | bilateral temporal lobe, right frontal lobe, cingulate gyrus & insula | right temporal lobe | right Anterior Temporal Lobectomy | moderate hypomnesia, anxiety | FCD Type IIa (Focal Cortical Dysplasia) |

### Experimental design and statistical analyses

#### Experimental paradigm

Participants performed a passive navigation task in a Virtual Reality (VR) environment within a circular track at a constant speed. Patients performed the task on a laptop while sitting in a comfortable position in their hospital bed at a distance of approximately 50 cm from the screen. The VR application was created using the Unity3D game engine (Unity Technologies, San Francisco, CA, USA). Four 3D models of buildings placed on the outer part of the track served as global environmental landmarks during navigation. Within the track, a set of local cues composed of everyday-life objects, obtained from the “texture city prefabs” library for Unity 3D (https://unity.com/) were randomly selected and placed at uniform distances along the navigational path (see Figure 1A for a bird-view of the virtual environment). The participants viewed the track from a first-person perspective. The starting point position in each trial was randomized but the order of items stayed the same and participants always faced the same direction. 60 trials in total were completed by each patient, lasting around 40 minutes.

**Figure 1.**
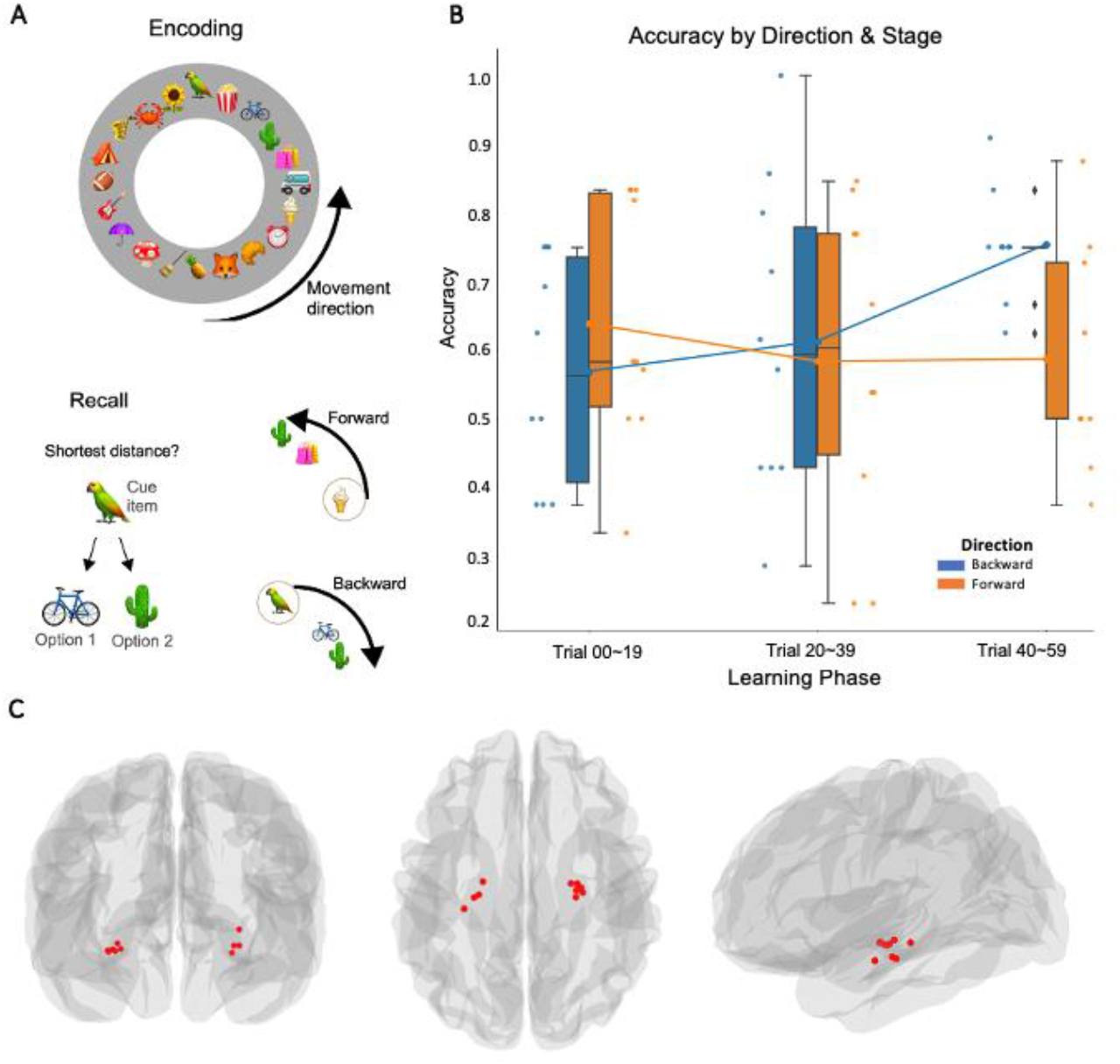
A: Illustration of the virtual navigation task paradigm. *Top:* bird-view layout of the circular navigation path (not shown to participants throughout encoding). *Bottom:* Example of a cued-distance judgment task during recall. Experimental condition: recall directionality (forward/backward), depending on the order in which items were encountered during passive navigation. B: Memory task accuracy, each dot represents a single participant’s performance at early, middle or late stage of the task, under forward or backward condition. Boxes denote the 25^th^ to 75^th^ percentile and the median line. Whiskers extend 1.5 times the interquartile range from the edges of the box. The lines on the boxplots indicate the means for each of the conditions. C: Electrode locations in the hippocampus from individual patients (n=10) in MNI space coordinates by front, top, and lateral views.

#### Navigation and memory task

Each trial started with a self-paced blank screen with a black fixation cross in the center alerting the subject of the beginning of a new trial. At each trial, the number of local landmarks placed in the surroundings of the circular track and the locomotion speed of the virtual avatar varied accordingly with the experimental condition. The number of visual cues placed on the track was set to either 50, 100, or 150 landmarks (determining the diameter of the circular track), and the speed of locomotion was set as 19.9, 26.7, or 32.7 deg/sec, for the low, medium, and high information density conditions respectively. Each navigational lap contains 20 items and the mean and standard deviation of the number of laps per trial are 1.43 ± 0.34 (see multimedia video). Details for the encoding stage and part of the related data were reported elsewhere (Santos-Pata et al., 2022).

At the end of the navigation, a blank screen appeared for 4 s, participants were then probed with a cue item and asked to indicate which of the two items was closer to the cue during navigation. For the memory judgment of relative temporal order, we operationally defined 2 levels of this condition by the order in which the cue and choices were encountered: forward or backward search (Figure 1A). Note that these 2 option items were always in the same direction to the cue, within 1 quadrant of the circular track (i.e., < 5 items apart). The trial presentation order was randomized across participants. Our analysis focused on the memory stage.

#### Electrophysiological recordings and electrodes localization

The location of the electrodes was established only for clinical reasons using an sEEG approach. Targeted regions varied across patients depending on their clinical assessment, but in all participants recruited for this study, they included the hippocampus in the left (n=4) or the right hemisphere (n=6). After co-registering pre- and post-electrode placement using MR scans and CT whole-brain volumes, we confirmed 10 pairs of contacts located in the hippocampus (Figure 1C). Locations of the electrodes in native space were finally converted to the standard Montreal Neurological Institute (MNI) space using 3D Slicer (https://www.slicer.org) and BrainX3. Recordings were made using a standard clinical EEG system (EEG-1200C, Nihon Kohden, Co.) at a sampling rate of 2000 Hz. A unilateral implantation was performed in all patients, with each of them having 8 to 11 intracranial electrodes (Sinovation, Beijing, diameter: 0.8 mm, 8 to 16 contacts, 2 mm length, 1.5 mm spacing between contact centers) stereotactically inserted with robotic guidance (Sinovation, Medical Technology, Ltd., Beijing, China).

#### Event related field potentials and preprocessing

When more than one contact was available, the electrode with higher delta-theta power ratio was selected. A bipolar montage was used offline to mitigate any effect of volume conduction or any confounds from the common reference signal. Bipolar signals were derived by differentiating electrodes pairs, the contact of interest, and one contact from adjacent white matter identified anatomically, of recorded and not rejected channels within the same electrode array. The continuous iEEG signals at the selected electrodes were first band-pass filtered between 1Hz and 200Hz using a two-way, zero phase-lag, finite impulse response filter to prevent phase distortion (*mne*.*filter*.*filter_data* function). To remove power line contamination, we applied a notch-filter at 50Hz and harmonics with a 2 Hz bandwidth (*mne*.*filter*.*notch_filter* function). Stimulus-triggered TTL pulses were also recorded with the iEEG data for synchronization with task events. For each patient, the hippocampal event related field potentials were computed by averaging the pre-processed signals from the selected contact points locked to the memory retrieval stage onset.

#### Power spectrum fit and peak frequency analysis

The power spectrum density was computed using the Welch method (using the *scipy*.*signals*.*welch* function) and fitted to the 1/f background spectrum in a log-log space to show frequency dependence of bands during recall task where oscillatory power exceeded 1 standard deviation over the 1/f background. We also calculated the number of oscillatory activities whose estimated peak frequency falls in each 1-Hz bin (1-15 Hz) using the FOOOF library (http://www.fooof-tools.github.io), and detected dominant frequencies during memory retrieval for each subject through permutation tests (n=1000) for later analysis.

#### P_episode_ *rate and analysis*

We utilized the oscillatory episode detection algorithm to identify high-power rhythmic activity. We used the measure P_episode_, which is defined as the proportion of time during which oscillations at a given frequency were present (Caplan and Glaholt, 2007, Caplan et al., 2001, Caplan et al., 2003, van Vugt et al., 2007; Whitten et al., 2011). We operationally defined an oscillatory episode at a frequency of interest, f*, as a duration that exceeded a time threshold, DT, during which power at frequency f* was greater than a power threshold, PT. We selected the values of PT and DT as follows. We applied a Morlet wavelet transform to the raw data traces to move to the frequency domain. We used 28 frequency steps logarithmically sampled in the range of 1-76 Hz to preserve the relative bandwidth. The wavelet transform provides us with wavelet power as a function of time at each frequency of interest. To determine the value of PT at each recording site for each frequency, we assumed that the background spectrum was “colored noise”, with the form of Af-α. We fitted the theoretical Power (f) =Af-α function to the actual power spectrum over the 1-to 76-Hz range at each electrode. We then took the fit value at the frequency of interest, f*, as the mean of the χ2(2) distribution of wavelet power at that frequency and set PT at that frequency to the 95^th^ percentile of the cumulative distribution of this fit χ2(2) function to exclude 95% of the background signal. The duration threshold, DT, at frequency f* was set to three cycles [i.e., 3(1/f*)] to eliminate artifacts and physiological signatures that were nonrhythmic. P episodes(f) were originally defined as the total time during which episodes occurred at frequency f divided by the total trial time. We adapted the measurement of P episode(f) by calculating the number of oscillatory events per second across the frequency bands of interest (2-8 Hz for theta and 30-70 Hz for gamma), namely the rate of P-episode.

#### Generalized linear mixed models

We built 2 linear mixed models with package lme4 (Bates, Maechler & Bolker, 2012) in R (R Core Team, 2012) analyzing the effect of directionality condition and stage of the task on rates of P-episode for theta and gamma oscillations respectively. Behaviorally, we ran 2 separate generalized linear mixed models with package car (Fox & Weisberg, 2019) to analyze the effect of directionality condition and stage of the task on response accuracy (binomial family, logit link function) and reaction time (inverse Gaussian family, inverse link function). For all models, we adopted a 2 × 3 design where 2 levels of recall directionality (forward vs backward) and 3 levels of learning phase (early vs middle vs late stage of the task) were entered as fixed effects, with intercept for subjects entered as random effects to account for individual differences in overall task performance and missing data points. Type 3 likelihood ratio tests comparing the full models against corresponding null models were used to obtain p-values.

## Results

### Memory performance

In terms of response accuracy, a significant interaction between the two fixed effects was found (*F (df=2) = 6*.*26, p =* .*044*), highlighting a diverging pattern of change in performance across stages of the task by recall directionality condition. Estimated marginal means (EM) with 95% confidence interval further suggested that at early stage of the task, response accuracy on forward trials (EM = 0.644, 95%CI = [0.540, 0.735]) was higher than backward trials (EM = 0.577, 95%CI = [0.459, 0.686]), however at late stage of the task, response accuracy on backward trials (EM = 0.773, 95%CI = [0.671, 0.851]) became higher than forward trials (EM = 0.598, 95%CI = [0.475, 0.709]) (Figure 1B). No significant effect of interaction was found on reaction time (*F (df=2) = 1*.*17, p =* .*558*). There was however a significant fixed effect of recall directionality (*F (df=1) = 6*.*49, p =* .*011*), where backward trials were faster than forward trials, as well as a fixed effect of task stage (*F (df=2) = 9*.*22, p =* .*010*), where responses were slower towards the end of the experiment.

### Frequency dependence during recall task

Before looking into the effects of experimental manipulations, we first demonstrated the frequency dependence of the recall task across subjects. By fitting the power spectrum density to the 1/f background spectrum in a log-log space, we identified the frequency bands where oscillatory power exceeded 1 standard deviation over the 1/f background (Figure 2 left, gray regions). Moreover, we calculated the number of oscillatory activities and detected subject-specific dominant frequencies (Figure 2 right, histograms). Both measures consistently revealed prominent oscillations during recall at theta band and low gamma band, which gave us an understanding of the frequency dependence for the task. After taking into account each subject’s variation, this result guided us to choose the frequency band of 2 - 8Hz (theta) and 30 - 70Hz (gamma) for the P_episode_ analysis.

**Figure 2.**
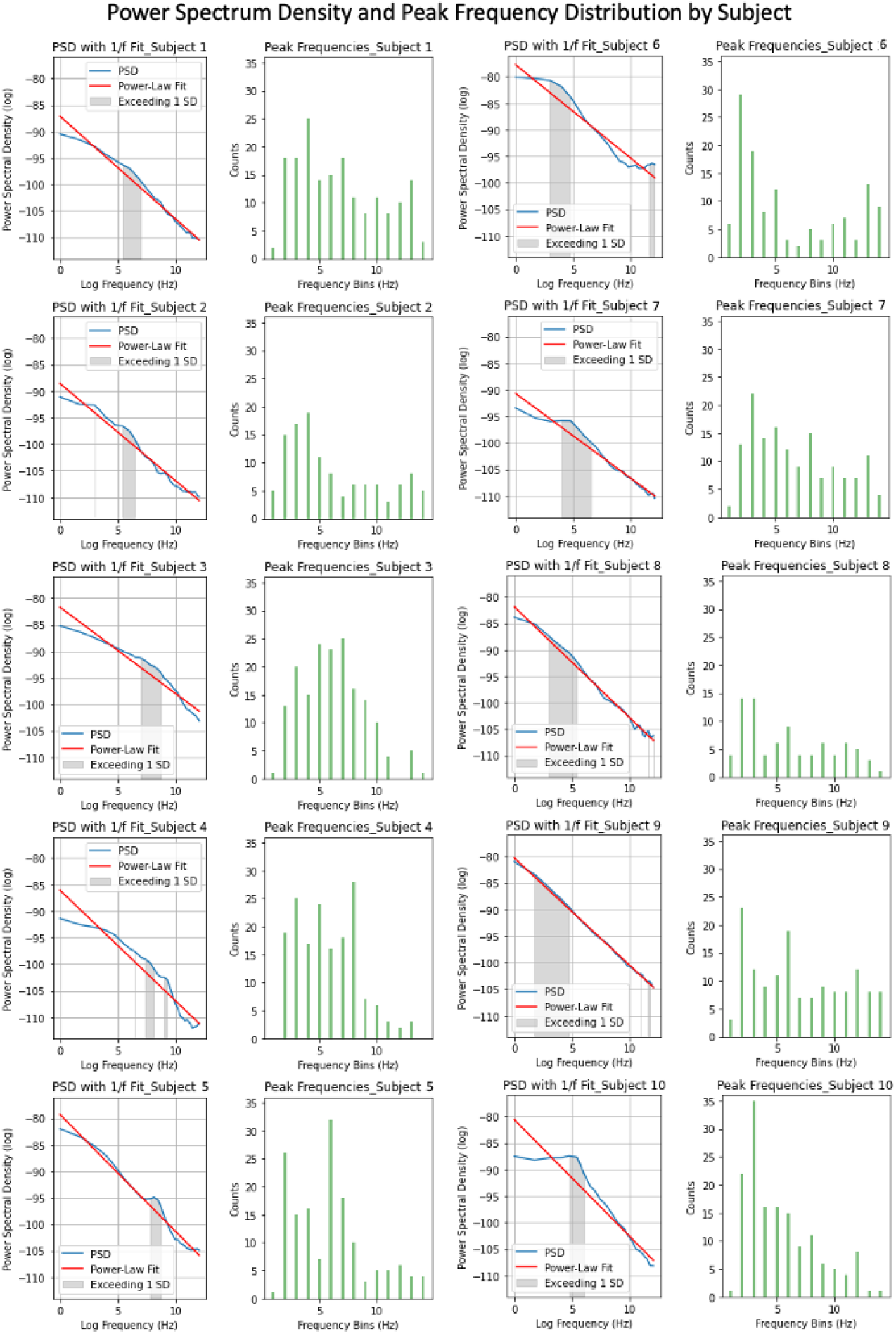
Power Spectrum Density for each subject in a log-log space, fitting to the 1/f background spectrum, with gray regions indicating frequency bands of power higher than 1 standard deviation of the 1/f background spectrum (left plots). Distributions of peak frequencies detected in each oscillatory activity are shown by histograms (right plots).

### P-episode rates

In line with memory accuracy, a significant interaction effect (*F (df=2) = 6*.*90, p =* .*032*) was revealed where the rates of theta P_episode_ during forward trials (EM = 0.374, 95%CI = [0.266, 0.482]) was higher than during backward trials (EM = 0.310, 95%CI = [0.200, 0.421]) at the early stage of the task, followed by a reversed pattern at the late stage of the task when the rates of theta P_episode_ became higher for backward trials (EM = 0.418, 95%CI = [0.308, 0.527]) than during forward trials (EM = 0.369, 95%CI = [0.258, 0.480]) (Figure 3A). For comparison, we performed a corresponding set of p-episodes analysis on the gamma band (30 - 70 Hz). We found no significant fixed effect of task stage or interaction but only a fixed effect on recall directionality P_episode_ (*F (df=1) = 5*.*59, p =* .*018*), where P_episode_ was higher for forward trials (EM = 0.595, 95%CI = [0.362, 0.828]) than backward trials (EM = 0.538, 95%CI = [0.304, 0.771]) (Figure 3B). The exact time points at which all the theta and gamma P_episode_ occurred for each trial by conditions are shown per patient (Figure 4).

**Figure 3.**
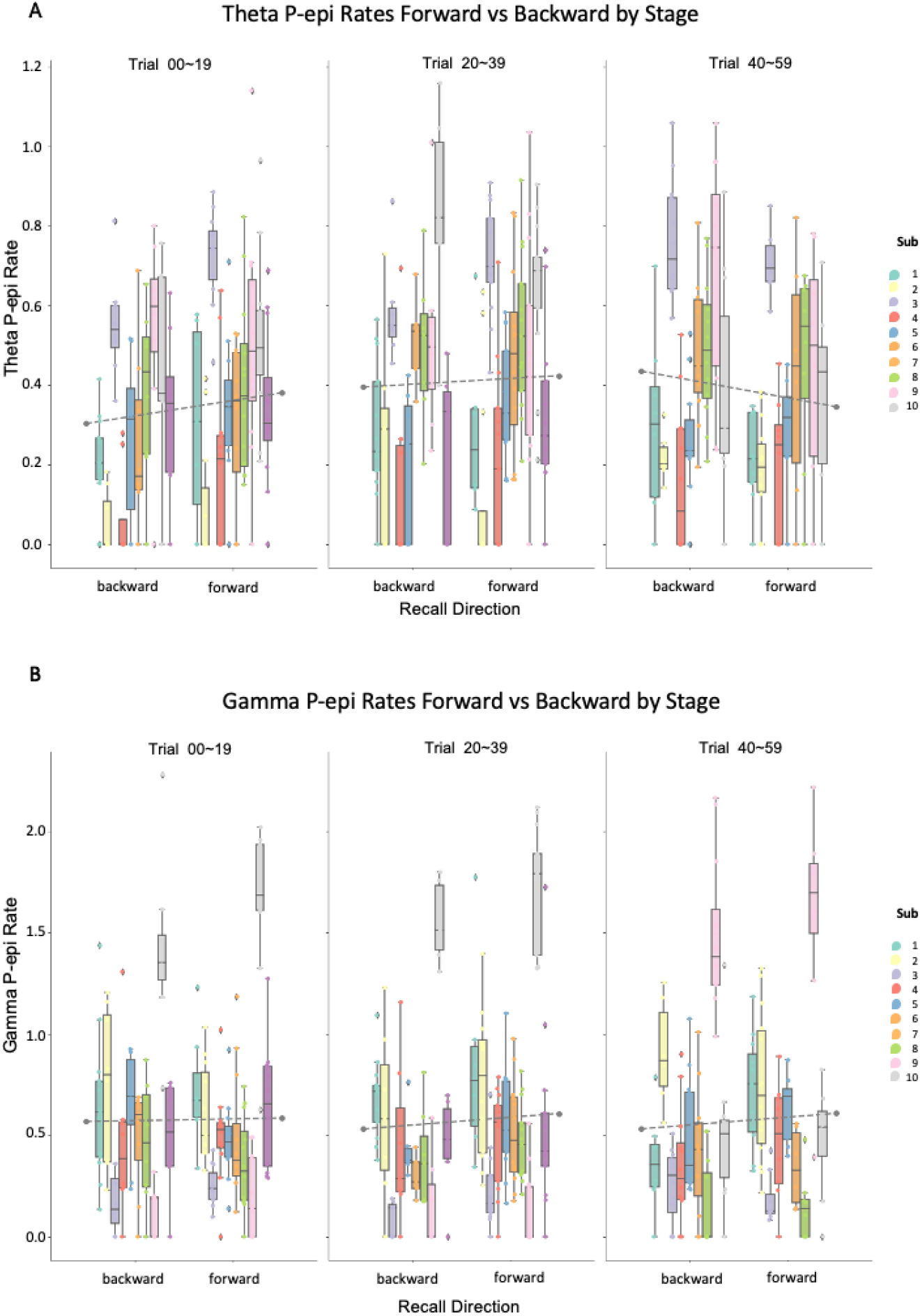
A: Theta P_episode_ rates organized by directionality and stage conditions. There is a directionality by stage interaction. B: Gamma P_episode_ rates organized by directionality and stage conditions. There is a main effect of directionality irrespective of stage. Each color denotes an individual patient. Boxes denote the 25^th^ to 75^th^ percentile and the median line. Whiskers extend 1.5 times the interquartile range from the edges of the box. Dashed lines on the boxplots indicate the mean across patients for the conditions.

**Figure 4.**
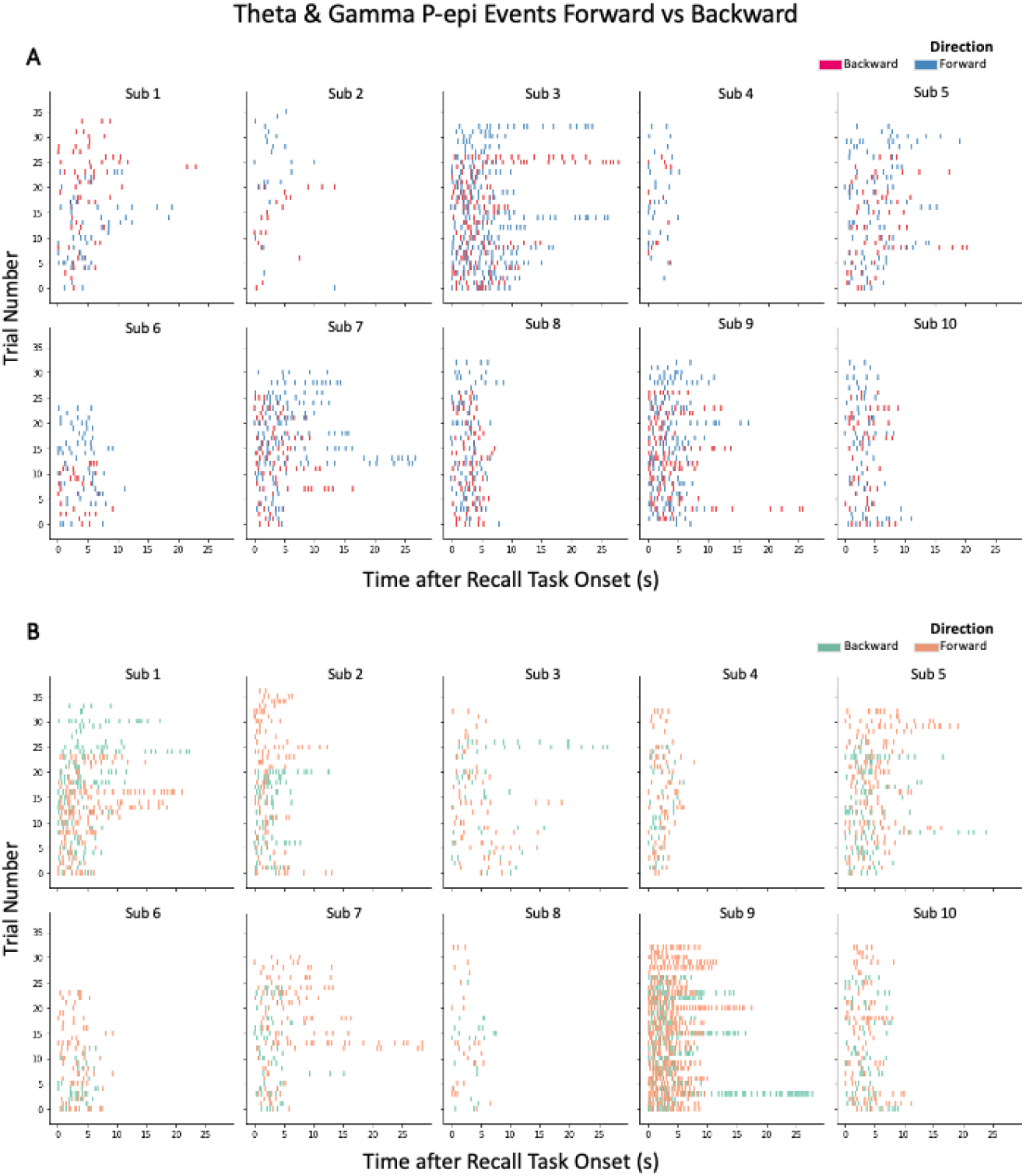
A: Theta P_episode_ events organized by trials per patient. B: Gamma P_episode_ events organized by trials per patient. Time zero is time-locked to memory test onset. Dots are color coded by retrieval directionality (forward/backward); each dot represents a single P_episode_ event.

## Discussion

In this study, the role of hippocampal theta activity in recall directionality was investigated using a relative order judgment task following encoding in a virtual circular track. The results showed that the sustained high-power oscillatory events in the theta band, quantified by P_episode_ rate, increased for the backward-cued than forward-cued recall conditions during the later stage, compared to the earlier stage of the task. This suggests that learning results in experience-dependent changes in the neural signal in the hippocampus. In the following we will discuss several contributions of the findings.

The hippocampus is known to code for abstract concepts and memories in some similar way as coding for space (Constantinescu et al, 2016; Bellmund et al, 2018; Solomon et al, 2019). This notion aligns with other recent work on hippocampal theta that are indicative of links between navigational space (Chen et al, 2021) and information search through other types of mental spaces, such as temporal and semantic clustering in memory recall (Sakon & Kahana, 2022; Sakon et al, 2022). The temporal order memory task here including the two spatial directionality is a hybrid of both abstracted space and memory representation. We demonstrated an interplay between theta oscillations and learning for temporal order memory processes. During initial stages of learning, theta events are biased towards forward retrieval in the cued-recall of forward/backward sequences. As learning occurs, the hippocampal theta episodes facilitate experience dependent changes for directionality in retrieval. This evidence on how human hippocampal theta P_episode_ is implicated in learning suggests an important link between theta oscillations and how they incorporate learning and experiences towards modulating our episodic memory representation and processing (Estefan et al, 2021). In comparison to the volitional learning reported in Pacheco Estefan et al. (2021), our experience-dependent learning is of an incidental nature. Another difference with Pacheco Estefan et al. is that their theta enhancement effects are manifested strictly in the encoding stage whereas we observed the change in theta power in the retrieval stage.

To ascertain that these putative effects are theta-specific, we performed a corresponding analysis using gamma oscillations and we found no such learning effects for gamma activity. We noted there was a main effect of directionality in gamma P_episode_ rates, where forward trials had consistently higher P_episode_ across all stages. In free call paradigms, there is a higher probability of successively recalling items in a forward vs. backward direction, where for a given recalled item, subsequently recalled items are more likely to have been encoded after (rather than before) the first-recalled item (Polyn & Kahana, 2008; Polyn, Norman, & Kahana, 2009). This forward asymmetry bias arises because a given item becomes part of the temporal context for succeeding items and serves as a memory cue. In our navigational case, a forward-cued condition, with it reinstating the sequence of items in an identical direction as encoding, would have reinstated more item representations within the sequence, thereby resulting in a strengthened temporal context upon subsequent memory test. Since gamma activity is involved in organizing and temporally segmenting the representations of different items (Howard et al, 2003; Pesaran et al, 2002), this might explain why gamma P_episode_ rates were consistently higher for forward than backward trials. The stability of gamma P_episode_ rates across all stages also suggests that the gamma correlates were not affected by the theta-mediated “learning” we observed here and is consistent with the initial performance advantage before any theta-mediated “learning” occurred (see Liu & Caplan, 2022; Liu, Chan & Caplan, 2014; Yang et al, 2013; Chan et al, 2009). In addition, inspired by studies on recall, as suggested recently by Dougherty et al. (2023), the dynamics of forward vs. backward memory processes are yet to be considered more thoroughly in extant memory models. Our findings on the differential involvements of theta and gamma P_episode_ might help inform the neural basis of current Retrieved Context Theories.

As evidenced by earlier studies, the hippocampal system is critical to enable normal learning of temporal structure (Kwok et al, 2015), by bridging spatial information across delay (Naya & Suzuki, 2011; Sakon et al, 2014) and/or accumulating a history of spatial relationships (Forcelli et al, 2014; Rueckemann & Buffalo, 2017). To relate our findings with the broader animal literature, our results echo with models stipulating a common neural substrate underlying relational encoding encompassing both memory and navigation (Howard & Eichenbaum, 2015). The discovery of cells sensitive to both space and time has also supported the relational memory system in its physiological plausibility (Kraus et al, 2013). Indeed, there has been evidence suggesting that a gradual change in the pattern of hippocampal activity would form a temporal context for events and support subsequent memory of the learned order (Manns et al. 2007). In light of other observations that hippocampal place fields are positively skewed in their initial states, before gaining a negatively skewed shape through learning or experiences in both rodents (Mehta et al, 2000) and humans (Reddy et al, 2021; Rutishauser et al, 2010), our finding extended this possibility to the memory retrieval stage as well as in a more abstracted cognitive space.

All in all, we showed that hippocampal theta and gamma activity are respectively involved in the retrieval of relational structures embedding temporal order memories. Specifically, the sustained theta oscillations reflect the learning processes obtained from repeated experiences. This study helps elucidate part of the electrophysiological mechanisms underlying the interplay between learning and spatiotemporal sequence memories upon retrieval in the hippocampus.

## AUTHOR CONTRIBUTIONS

Designed research: HJ, JC, DSP, SCK; Performed research: HJ, JC, DSP, JT, YC, JY, CL, XZ, RW, LS, SZ; Contributed unpublished reagents/analytic tools: DSP, JY, CL, LS, SZ; Analyzed data: JC, DSP; Wrote the paper: JC, DSP, SCK

## ACKNOWLEDGMENTS

The research results of this publication are sponsored by the Kunshan Municipal Government research funding Project Number: 23KKSGR017.

## DATA AND CODE AVAILABILITY

Raw electrophysiological data, analysis code, and processed data supporting the conclusion of this study are available upon request.

